# Chlorhexidine reduced susceptibility associated to tetracycline resistance in clinical isolates of *Escherichia coli*

**DOI:** 10.1101/2021.10.05.463149

**Authors:** Guilhem Royer, Jose-Manuel Ortiz de la Rosa, Xavier Vuillemin, Béatrice Lacombe, Françoise Chau, Olivier Clermont, Mélanie Mercier-Darty, Jean-Winoc Decousser, Jean-Damien Ricard, Patrice Nordmann, Erick Denamur, Laurent Poirel

## Abstract

Chlorhexidine is a widely used antiseptic in hospital and community healthcare. Decreased susceptibility to this compound has been recently described in *Klebsiella pneumoniae* and *Pseudomonas aeruginosa*, together with cross-resistance to colistin. Surprisingly, few data are available for *Escherichia coli*, the main species responsible for community and healthcare-associated infections. In order to decipher chlorhexidine resistance mechanisms in *E. coli*, we studied both *in vitro* derived and clinical isolates through whole-genome sequence analysis. Comparison of strains grown *in vitro* under chlorhexidine pressure identified mutations in the gene *mlaA* coding for a phospholipid transport system. Phenotypic analyses of single-gene mutant from the Keio collection confirmed the role of this mutation in the decreased susceptibility to chlorhexidine. However, mutations in *mlaA* were not found in isolates from large clinical collections. In contrast, genome wide association studies (GWAS) showed that, in clinical strains, chlorhexidine reduced susceptibility was associated with the presence of *tetA* genes of class B coding for efflux pumps and located in a Tn*10* transposon. Construction of recombinant strains in *E. coli* K-12 confirmed the role of *tetA* determinant in acquired resistance to both chlorhexidine and tetracycline. Our results reveal two different evolutionary paths leading to chlorhexidine decreased susceptibility: one restricted to *in vitro* evolution conditions and involving a retrograde phospholipid transport system; the other observed in clinical isolates associated with efflux pump TetA. None of these mechanisms provides cross-resistance to colistin or to the cationic surfactant octenidine. This work demonstrates the GWAS power to identify new resistance mechanisms in bacterial species.

## Introduction

Chlorhexidine (CHX) is a widely used antiseptic in hospital settings as well as a community-based care (oral hygiene, wound antiseptic, antiseptic scrubs, etc…) (1). In the past few years, some authors reported reduced susceptibility to this cationic agent in *Klebsiella pneumoniae* (2) and *Pseudomonas aeruginosa* (3). They also reported cross-resistance to colistin (COL), a polypeptide antibiotic with activity against most Gram-negative bacteria (4). Such cross-resistance between one of the most frequently used antiseptic and a last resort antibiotic is of great concern in the context of ever-growing antimicrobial resistance (5). However, this CHX-COL cross-resistance was mainly observed from *in vitro* obtained mutants and it is not clear to what extent they can be transferred to clinical microbiology (2, 3, 6–8). Moreover, only few data are available regarding that concern in *Escherichia coli*, the main species responsible both for community and healthcare-acquired infections (https://www.ecdc.europa.eu/en/healthcare-associated-infections-acute-care-hospitals/database/microorganisms-and-antimicrobial-resistance/most-frequent). Of note, Gregorchuk *et al.* (9) recently identified a limited set of determinants potentially involved in CHX reduced susceptibility in E. coli using a multi-omics approach. By contrast to what had been observed with the other species studied, cross-resistance with COL was not observed in that study. These data were obtained from experimental *in vitro* obtained mutants of *E. coli* K-12 BW25113 (10).

To take into account the potential impact of the ecological niche, we recently compared the distribution of COL MICs and maximum growth rate under CHX 2 mg/L from two collections of *E. coli* clinical isolates, *i.e.* i) the ColiRed collection including 153 COL-resistant fecal strains (11) and ii) the ColoColi collection which consists of 215 pulmonary strains characterized by a high frequency of reduced susceptibility to CHX (12). From this comparison, no positive correlation could be identified between the colistin and the antiseptic resistance, suggesting that this cross-resistance was rare among clinical isolates (13).

In the present study, the molecular bases of mechanisms leading to decreased susceptibility to CHX were determined, using both *in vitro* obtained mutants and also clinical isolates. To this end, we analyzed CHX-resistant strains obtained after *in vitro* serial passages with CHX as well as from collections of clinical isolates. By comparing the different genomes of the corresponding strains, two different evolutionary paths leading to decreased susceptibility to CHX were identified.

## Material and methods

### Bacterial strains

Clinical isolates analyzed in the present study had been recovered from two previously published collections, namely ColiRed (11) (bioproject PRJEB28020) and ColoColi (12) (bioproject PRJEB39268). Briefly, the ColiRed collection consists of 153 COL-resistant *E. coli* strains isolated from rectal swabs of patients hospitalized in six university hospitals in the Paris area (“Assistance Publique-Hôpitaux de Paris”) between January and March 2017. The ColoColi collection is made up of 215 *E. coli* pulmonary strains retrieved from mechanically ventilated patients in French intensives care units and characterized by a high proportion of decreased susceptibility to CHX.

From the ColoColi collection, we selected three CHX-susceptible isolates from three distinct phylogroups that we submitted to CHX selection pressure: AN03, AN43, RY10. None of them carried class B *tetA* genes. These strains were serially passed with various concentrations of CHX. Briefly, after 18h incubation in lysogeny broth (LB), culture was diluted 1/1000. Then, 225 μL of this suspension were added to 25μL of a CHX solution with concentrations ranging from 5 to 1,280 mg/L in 96-well plates (Corning Inc., NY, USA) and incubated at 37°C. Minimum inhibitory concentrations (MICs) were read at 24 h of incubation and the same protocol was applied again to the well that grew with the highest CHX concentration (*i.e.*, the sub-inhibitory concentration). A total of eight serial passages were performed for each strain. From this experiment, three strains presenting elevated MICs of CHX, further defined as “resistant” (namely AN03CHX, AN43CHX and RY10CHX), were obtained.

To confirm the role of specific genetic determinant(s) involved in CHX decreased susceptibility, isolates from the Keio collection (10) were also investigated. Hence, the parental strain *E. coli* K-12 BW25113 was compared with the single deleted mutant lacking the gene of interest, namely JW2343 (Δ*mlaA*) and JW1771 (Δ*mipA*).

Finally, the *E. coli* K-12 derivative strain TOP10 (Invitrogen, Fischer Scientist) was used to further construct transformants with candidate genes involved in CHX resistance (see below).

### Susceptibility testing

Reduced susceptibility to CHX was assessed by measuring bacterial growth in LB supplemented with 2 mg/L CHX over a 24h period with an automatic reading of optical density (Tecan Infinite F200 pro) at 600 nm as previously described (La Combe et al., 2018). Strains with a maximum growth rate (MGR) > 0.8 h^−1^ were considered to have a reduced susceptibility to CHX (La Combe et al., 2018). We also performed the same experiment on a subset of strains with 1 mg/L CHX to identify low level decreased susceptibility, using the same threshold of MGR > 0.8 h^−1^. MICs of COL, octenidine, a novel antiseptic not structurally related to CHX (14), and tetracycline were determined by the broth microdilution method according to the EUCAST recommendations.

### Genomic comparison of *in vitro* evolved isolates and Keio collection strain analysis

The genomes of each strain (AN03, AN43, RY10) as well as their most resistant evolved counterpart (AN03CHX, AN43CHX, RY10CHX) were sequenced using Illumina sequencing technology, as previously described (11). Genomes were assembled with shovill v1.0.4 (https://github.com/tseemann/shovill) and annotated with Prokka v1.14.5 (15). The phylogroups of each strain were determined using the ClermonTyping (16) and their Sequence Type (ST) according to the Warwick scheme (17). We searched for single nucleotide polymorphisms and insertion/deletion (SNP/indel) using breseq v0.33.1 (18). SNPs detected as false positive when comparing mapping reads of a given strain to the assembly of the same strain were discarded. Thereafter, reads from each resistant strain were mapped onto the assembly of the corresponding susceptible one. Functional effect of the non-synonymous mutations was predicted using Provean (19), Polyphen-2 (20) and SIFT (21), and interpreted as previously described (22) (*i.e.* damaging if predicted by at least two softwares). For genes in which truncating mutations were identified, the MGR CHX 2 mg/L of the single-gene deleted corresponding strain from the Keio collection (JW2343 and JW1771) was further measured (10), using K-12 BW25113 strain as control. MIC of COL was also measured for the three pairs of strains as well as the strains from the Keio collection.

### Targeted approach to identify mutations responsible for CHX mutation in natural isolates

Putative mutations possibly responsible for reduced susceptibility to CHX were searched for among genome sequences recovered from clinical isolates belonging to both collections, namely ColiRed (11) and ColoColi (23). We performed a targeted approach by focusing on genes identified as potential resistance determinants through the comparison of *in vitro* evolved strains (seeking for differences between the wild-type and evolved strain counterparts, respectively). Additionally, we focused our search on genes encoding efflux, porins and global regulators of antibiotic resistance, as previously described (22) (Table S1). Meanwhile, we computed the pangenome of all strains and performed an alignment of the core genes using PPanGGOLiN (24). From this alignment, a maximum likelihood phylogenetic tree was computed using Iqtree v1.6.12, as previously described (25, 26). This phylogenetic tree was used to differentiate polymorphisms from candidate mutations, *i.e.* mutations that are found multiple times in resistant strains.

### Genome wide association study (GWAS) to identify without a priori mechanisms of reduced susceptibility to CHX

In a more exploratory way, a non-targeted approach was performed, consisting in a GWAS based on the gene presence/absence matrix previously obtained with PPanGGOLiN. Scoary was used with standard parameters and MGR CHX 2 mg/L > 0.8 h^−1^ as phenotype of interest, with the goal to potentially identify genes / gene families associated with decreased susceptibility to CHX (27). Only hits with a Bonferroni corrected p-value <0.05 were further considered. The genomic localization of the potential genetic determinants identified using blastN was analyzed against the nucleotide collection of NCBI database (https://blast.ncbi.nlm.nih.gov/Blast.cgi).

### Plasmid constructs

To test the candidate genes identified through GWAS, two recombinant strains were produced, using *E. coli* TOP10 as recipient strain (Invitrogen, Fischer Scientist) and pTOPO as plasmid vector, with corresponding inserts respectively containing the whole transposon *tetR *tetA* tetC* or the *tetA tetC* genes only. Those recombinant strains were constructed using purified DNA of a *TetA*-positive *K. pneumoniae* clinical isolate from our lab collection as template. The insert corresponding to the whole tetR-tetA-tetC operon was amplified using primers TetR-R (5’-GGT GGT TAA CTC GAC ATC TT-3’) and TetR (5’-GCG AGA ATG CTG TTC AAT AT-3’) and the tetA-tetC insert using primers TetF (5’-ACC ACT CCC TAT CAG TGA TA-3’) and TetR. After electro-transformation of the recombinant plasmid into *E. coli* TOP10, selection of the recombinant strains was performed using trypticase soy agar plates supplemented with tetracycline 8 μg/ml.

### Statistics

A Khi2 test was performed to compare the proportion of resistant strains according to their tetA status, with a threshold of significance fixed at 0.05. A Student t-test was performed to compare MGR in LB without CHX for JW2343 and the wildtype strain BW25113.

## Results

### *in vitro* selection of CHX reduced susceptibility associated with modification of the regulation of phospholipid transport system coding gene *mlaA*

Three CHX susceptible strains (AN03, AN43 and RY10) were selected from the ColoColi collection. They were used for selection on increased concentrations of CHX by serial passages (AN03CHX, AN43CHX and RY10CHX). After eight passages, MGR CHX 2 mg/L was > 0.8 h^−1^ for the three evolved strains. No effect was observed on MICs of COL for those different mutants. The genomes of those mutants were then sequenced to further perform SNPindel analysis. Genomic comparison of pairs of resistant and susceptible strains allowed to identify very few single nucleotide polymorphisms (SNPs) responsible for major protein variation (Table 1). Either a deletion of an adenine nucleotide being subsequently responsible for a frameshift or a nucleotide substitution leading to a premature stop codon were observed in the *mlaA* gene encoding MlaA (intermembrane phospholipid transport system lipoprotein) in isolates AN03CHX and RY10CHX, respectively. In addition, a missense mutation was also observed in the *barA* gene in strain AN03CHX being predicted to modify protein function. In the third strain, namely AN43CHX, a SNP responsible for a premature stop codon in the gene *mipA* encoding an MltA-interacting protein was identified, as well as a missense mutation in the *mscK* gene encoding a mechanosensitive channel with a probable effect on the protein function. Moreover, a detailed analysis of the genome sequences of strain AN43CHX identified the insertion of an IS*1*-like insertion sequence between *yfdC*, which code for a putative transport membrane protein, and *mlaA*. This insertion located 39 bp upstream of the *mlaA* start codon disrupted the transcription factor binding site (CACCTAAA) within the promoter of *mlaA* as predicted by BPROM (28), therefore likely interfering in the corresponding gene expression.

**Table 1.** Maximum growth rate, minimal inhibitory concentration of colistin and non-synonymous mutations identified in the pairs of **E. coli** strains evolved **in vitro** after 8 serial passages with chlorhexidine.

| Strains | Phylogroup | MGR CHX 2 mg/L (h<sup>-1</sup>) | MIC of colistin (mg/L) | Gene | Protein function | Nucleic variation | Protein variation | Type of mutation* |
| --- | --- | --- | --- | --- | --- | --- | --- | --- |
| AN03 | G | 0.02 | ≤ 0.06 |  |  |  |  |  |
| AN03CHX | G | 1.33 | ≤ 0.06 | *mlaA* | Intermembrane phospholipid transport system lipoprotein | 107delA | N36TfsX11 | Truncating mutation |
|  |  |  |  | *barA* | Signal transduction histidine-protein kinase | 1457C>T & 1458G>T | L486P | Substitution (damaging) |
| AN43 | D | 0.03 | ≤ 0.06 |  |  |  |  |  |
| AN43CHX | D | 1.47 | ≤ 0.06 | *mipA* | MltA-interacting protein | 73A>T | K25X | Truncating mutation |
|  |  |  |  | *mscK* | Mechanosensitive channel | 1783T>A | F595I | Substitution (damaging) |
|  |  |  |  | intergenic region |  | IS |  | Gene expression? |
| RY10 | B2 | 0.02 | ≤ 0.06 |  |  |  |  |  |
| RY10CHX | B2 | 0.97 | ≤ 0.06 | *mlaA* | Intermembrane phospholipid transport system lipoprotein | 97G>T | E33X | Truncating mutation |
| BW25113 (K-12 Keio collection) | A | 0.02 | ≤ 0.06 |  |  |  |  |  |
| JW2343 (Δ*mlaA*) | A | 0.85 | ≤ 0.06 | *mlaA* | Intermembrane phospholipid transport system lipoprotein |  |  | Single-gene deletion |
| JW1771 (Δ*mipA*) | A | 0.01 | ≤ 0.06 | *mipA* | MltA-interacting protein |  |  | Single-gene deletion |

MGR: Maximum growth rate
CHX: Chlorhexidine
MIC: Minimum inhibitory concentration
\*Predicted effect of the mutation is provided (see Material and methods section)

As a result of these genotypic analyses, a focus was made on the *mlaA* and *mipA* genes. The MGR CHX 2 mg/L was measured for the single-gene deleted strains from the Keio collection and the parental K-12 strain, namely JW2343 (Δ*mlaA*), JW1771 (Δ*mipA*), BW25113 (K-12 parental strain). Only JW2343 showed a high MGR, while strains JW1771 and BW25113 remained highly susceptible to CHX (Table 1). Hence, a convergence (the occurrence of distinct mutations in the same gene of phylogenetically unrelated strains) was observed in the *mlaA* gene under exposure *in-vitro* to CHX, corresponding to a strong selection signal (29), being therefore a strong indicator that the product of *mlaA* is involved in reduced susceptibility to CHX in *E. coli*. No fitness cost was observed when the cells were grown in a rich culture medium as LB [median MGR = 1.22 h^−1^ for the mutant JW2343 versus 1.29 h^−1^ for the WT (BW25113) strain, t-test *p-value*=0.37].

### Mutations in *mlaA* or in known efflux systems or porins do not play a critical role in the reduced susceptibility to CHX in clinical isolates

Then, a screening was performed based on the genome sequences available from the collections of clinical isolates previously mentioned. Among these 368 strains, 25 strains showed reduced susceptibility to CHX according to their MGR CHX 2 mg/L (13). First, we used a knowledge-based targeted approach and focused on genes previously identified as mutated through *in vitro* evolution experiments, namely *mlaA, mipA, barA* and *mscK*. By performing this targeted analysis, none of the mutations in those genes were found among clinical isolates (Table S2). In a second step, we analyzed the sequences of genes involved in efflux systems (n=15), encoding porins (n=5) or global regulator of resistance (n=9) (Table S1). A particular focus was made on those amino acid substitutions specifically identified among strains showing reduced susceptibility although being unrelated to phylogeny. Hence, a total of 16 substitutions were identified only in strains showing an MGR > 0.8 h^−1^. However, none of these substitutions was found more than one time among all those genome sequences (Table S2), ruling out any convergent mutation in those genes.

Consequently, those results strongly suggest that mutations in the *mlaA, mipA, barA*, and *mscK* genes, but also in genes coding for major efflux system and porins do not play a pivotal role in cross resistance to CHX.

### GWAS points to the role of efflux pump encoding class B *tetA* gene in reduced susceptibility to CHX

Given the results of the previous targeted approach, a more exploratory analysis was performed. From the pangenome data available from our strains, a GWAS was performed to uncover genetic determinants associated to reduced susceptibility to CHX. Interestingly, nine genes were found to be significantly associated with an MGR CHX 2 mg/L > 0.8 h^−1^ (Table 2). Among them, six genes were frequently associated with sequences specific for transposon Tn*10* (Genbank accession number: AY528506.1) carrying class B *tet*-related efflux genes, namely *tetA, tetR*, and *tetC*. These genes encode a tetracycline efflux MFS (Major Facilitator Superfamily) transporter, a transcriptional regulator and a tetracycline resistance-associated transcriptional repressor, respectively. Another *tet* gene, *tetD* encoding a transcriptional regulator, was also frequently associated (p-value = 0.08, close to the defined cut-off). The remaining three other genes for which a frequent association was found encoded poorly characterized proteins, being a bifunctional chitinase/lysozyme and two hypothetical proteins. These last three determinants were found in only five strains which all belong to the same sequence type (ST88, phylogroup C). Therefore, through this GWAS analysis, our hypothesis was that the *tet* operon might be likely involved in reduced susceptibility to CHX.

**Table 2.** Best hits (Bonferroni corrected p-value <0.05) from the GWAS analysis.

| Gene name | Protein annotation | Number of strains |  |  |  | Bonferroni corrected p-value |
| --- | --- | --- | --- | --- | --- | --- |
|  |  | MGR > 0.8 h-1 with the candidate gene | MGR < 0.8 h-1 with the candidate gene | MGR > 0.8 h-1 without the candidate gene | MGR < 0.8 h-1 without the candidate gene |  |
| unnamed gene | hypothetical protein | 15 | 23 | 10 | 320 | < 0.001 |
| *tetA* | tetracycline efflux MFS transporter class B | 16 | 33 | 9 | 310 | < 0.001 |
| *tetR* | TetR family transcriptional regulator | 16 | 33 | 9 | 310 | < 0.001 |
| *tetC* | tetracycline resistance-associated transcriptional repressor TetC | 15 | 29 | 10 | 314 | < 0.001 |
| unnamed gene | antibiotic biosynthesis monooxygenase | 10 | 7 | 15 | 336 | < 0.001 |
| unnamed gene | winged helix-turn-helix transcriptional regulator | 14 | 27 | 11 | 316 | < 0.001 |
| unnamed gene | bifunctional chitinase/lysozyme | 5 | 0 | 20 | 343 | 0,042 |
| unnamed gene | hypothetical protein | 5 | 0 | 20 | 343 | 0,042 |
| unnamed gene | hypothetical protein | 5 | 0 | 20 | 343 | 0,042 |

To test experimentally our findings, we constructed recombinant strains using *E. coli* TOP10 (K-12 derived) as the recipient and plasmid pTOPO as vector, giving rise to *E. coli* (pTOPO-*tetA-tetC*) and *E. coli* (pTOPO-*tetR-tetA-tetC*) respectively encompassing the corresponding genes (Figure 1A). Although the MIC of tetracycline was at 0.12 mg/L for *E. coli* TOP10 alone, it was measured at 16 mg/L for both *E. coli* (pTOPO-*tetA-tetC*) and *E. coli* (pTOPO-*tetR-tetA-tetC*). Then, the MGR of the *E. coli* recombinant strains was measured with increasing concentrations of CHX, using strains AN03 and AN03CHX as susceptible and resistant control strains, respectively (Figure 1B). Firstly, our data showed that *E. coli* TOP10 strain had a much lower fitness as compared to clinical isolate AN03 and its derivative AN03CHX when grown in absence of CHX, precluding the analysis of the impact of tet genes in concentrations of CHX equal or higher than 2 mg/L. Nevertheless, by measuring the MGR of *E. coli* TOP10 recombinant strains, a reduced susceptibility to CHX was found. Furthermore, since a cross resistance to CHX and COL has been suggested in other species, MICs of COL and octenidine were also measured for the recombinant strains, showing that the Tet efflux system did not have any impact on COL or on octenidine susceptibility.

**Figure 1:**
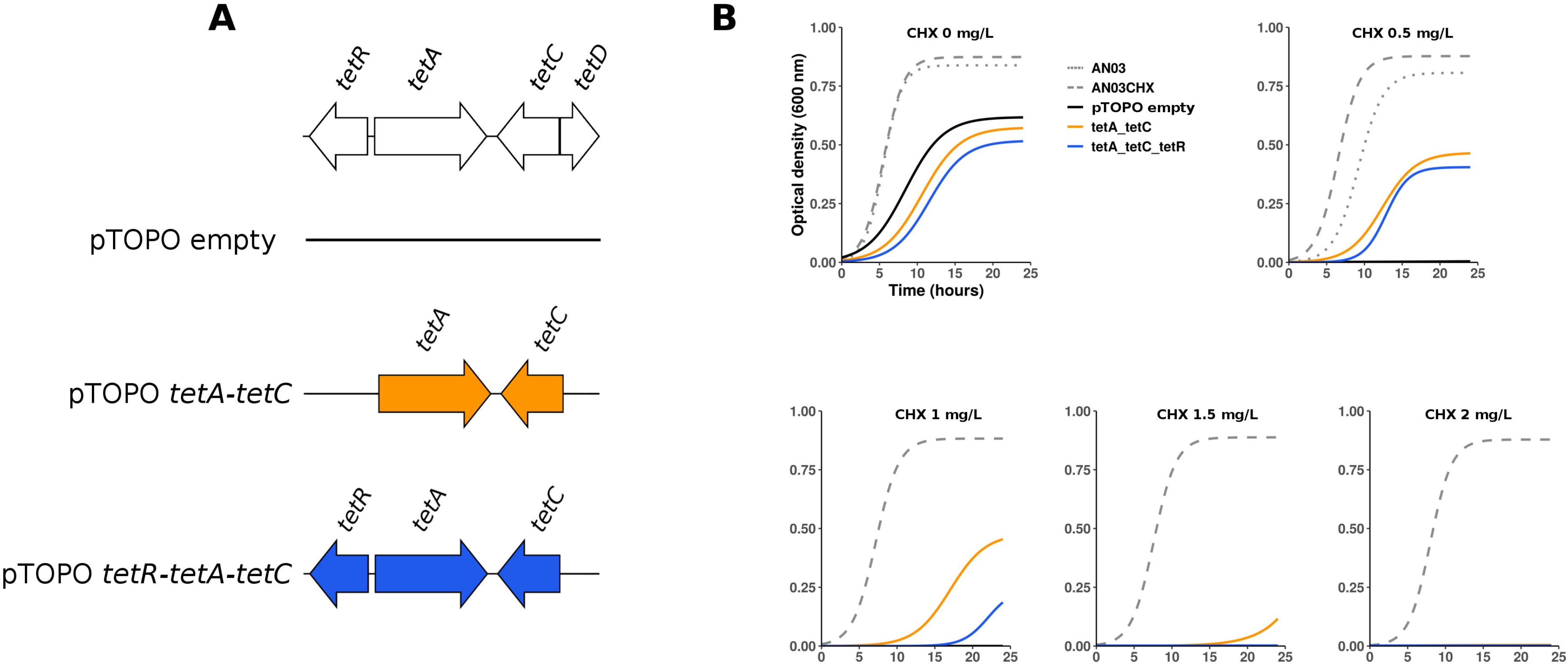
Analysis of the role of class B tet determinant in CHX decreased susceptibility. A) Transformant constructions based on the pTOPO platform. B) Growth curves with increasing concentration of CHX from 0 to 2 mg/L of the AN03 and AN03CHX strains used as negative and positive controls, respectively and the TOP10 strain with the different constructs described in A. Note the difference of growth rate without CHX between the natural isolate and its derivative on one hand and the TOP10 strain with the various pTOPO constructs on the other hand.

Consequently, GWAS allowed to identify the class B *tetA* gene as a source of reduced susceptibility to CHX. In other words, those tetracycline resistance determinants had a significant and collateral impact on susceptibility to CHX.

### Presence of class B *TetA* leads to low-level decreased susceptibility to CHX

Although an association between the *tetA* gene and a reduced susceptibility to CHX was identified, whole genome sequence analysis revealed a total of 33 isolates carrying *tetA* but exhibiting a MGR CHX 2 mg/L < 0.8 h^−1^. Consequently, we decided to evaluate the MGR of the *tetA*-positive strains with a lower CHX concentration (1 mg/L). A control group was used, being made of 48 *tetA*-negative strains, the choice of those strains being made by considering a significant match in terms of phylogroups between the control group and the group of strains to be tested. It showed that strains carrying *tetA* statistically had more frequently an MGR 1 mg/L > 0.8 h^−1^ (n=46/49 vs 11/48, p < 0.01). Moreover, by testing the three *tetA*-positive strains with an MGR 1 mg/L < 0.8 h^−1^, two had limit values (0.70 and 0.67 respectively). The third one had an MGR 1 mg/L of 0.06 h^−1^, but a detailed analysis of the sequence showed that the *tetA* gene was truncated by an IS*1* in that given isolate. Besides, genome sequence analysis did not evidence any key features that may explain reduced susceptibility to CHX among the eleven *tetA*-negative strains showing an MGR 1 mg/L > 0.8 h^−1^. Altogether, our data indicated that TetA of class B is involved in decreased susceptibility to CHX.

### The class B *tetA* gene is carried by genetically diverse strain backgrounds of *E. coli*

The 49 isolates that carried tetA of class B belonged to various phylogroups of *E. coli* (Figure 2), being respectively A (n=9), B1 (n=3), B2 (n=17), C (n=5), D (n=8), F (n=4), and to the cryptic clade I (n=1). Likewise, these isolates belonged to 31 different sequence types (STs) and to 26 O:H serotype combinations, highlighting their significant genetic diversity.

**Figure 2:**
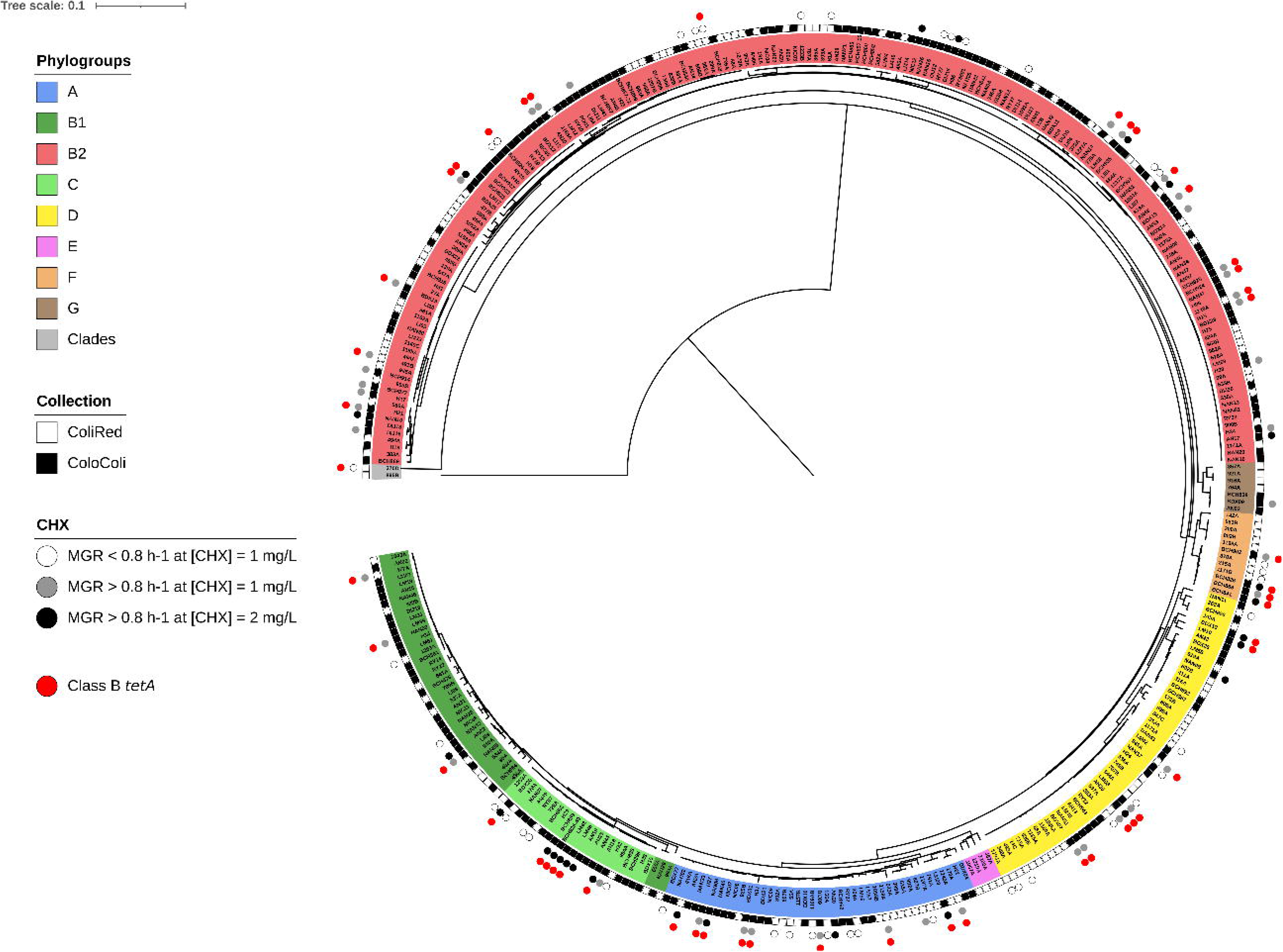
Core-genome based maximum likelihood phylogenetic tree of the 368 strains from ColiRed and ColoColi collections. Strains with decreased susceptibility to 1 mg/L and 2 mg/L of CHX are highlighted by grey and black points, respectively. Strains that exhibit an MGR < 0.8 h^−1^ in presence of 1 mg/L of CHX are highlighted by white points. The remaining strains did not have decreased susceptibility to 2 mg/L of CHX. The presence of class B tetA gene is indicated by red points. Strains are colored according to their genogroup. The source collection of the strains is indicated by white and black squares for ColiRed and Colocoli, respectively. The tree is rooted on the strain 895B which belongs to clade V. The scale represents genetic distance in nucleotide substitution per site.

## Discussion

Based on results of *in vitro* evolution experiments under CHX pressure, a convergent evolution could be identified with truncating mutations in the *mlaA* gene encoding an intermembrane phospholipid transport system lipoprotein. The pivotal role of this gene has been very recently confirmed by others by a multi-omic approach, including transcriptomic, proteomic and genomic analysis (9). As suggested in that latter study, the inactivation of the *mlaA* gene could likely lead to a lack of transport of CHX-phospholipid complex, as well as a lack of phospholipid recycling at the outer membrane. Moreover, the *mlaA* gene was also shown to be involved in hypervesiculation phenomenon (30). It can therefore be also hypothesized that these numerous outer membrane vesicles could act as a decoy likely catching a significant amount of the CHX.

Noteworthy, no cross resistance with COL could be observed by testing those strains. In fact, a CHX-mediated “cross-resistance” is still debated. Such phenomenon was reported in various *Enterobacteriaceae* and for several antibiotics after low level exposure to CHX, but this was observed only in isolates being selected *in vitro* (31). Likewise, no cross resistance was observed with octenidine, a molecule that shows excellent efficacy to eradicate multidrug-resistant Gram-negative and Gram-positive bacteria (14, 32).

Our analysis did not allow to correlate those mutations identified among *in vitro* obtained mutants with mechanisms responsible for CHX resistance among clinical isolates. In term of fitness, the inactivation of the *mlaA* gene through mutations likely induces a high burden owing to dramatic modifications of cell integrity as formerly demonstrated (9). Of note, such fitness cost might probably be occurring in an environment-dependent manner, as observed for other antimicrobial agents (33–35). Alterations of membrane phospholipids impair the adaptability to environmental stresses (36). Indeed, we did not evidence any fitness cost in the rich medium LB and the controlled experimental conditions of our evolution experiments are very far from the natural ecological niches of *E. coli*. A more comprehensive and targeted approach focusing on genes involved in efflux or coding for porins and global regulator of resistance also failed to identify determinants associated to CHX reduced susceptibility.

Nevertheless, our GWAS analysis allowed to get more insight into CHX resistance by identifying an association with the class B *tetA* gene encoding tetracycline resistance. This *tetA* gene belongs to the class B of *tet* genes, and is frequently found in *E. coli*. Indeed, from the recently published data on antibiotic resistance in EnteroBase (37), 12.5% of the *E. coli* genomes carry class B *tet* genes. By constructing recombinant *E. coli* strains, we could definitely assess the role of this determinant both in tetracycline resistance and CHX reduced susceptibility, the latter sometimes being moderate. Nevertheless, and as observed for the *in vitro* mutants, no impact on colistin resistance was observed for those strains carrying a *tetA* gene.

Interestingly, reduced susceptibility to CHX does not appear to be provided by all *tet* genes classes. Among our isolates, 96 carried at least one non-class B *tet* gene: *tetA* of class A (n=93), *tetA* of class C (n=1), *tetA* of class D (n=3), *tet* of class M (n=3). However, among these isolates only six had an MGR CHX 2 mg/L > 0.8 h^−1^ among which 4 also carried class B *tet* genes. Among the remaining two, one had a class D *tetA* gene which exhibits higher identity at the protein level with class B TetA than the other classes (Figure S1). Since class M genes do not encode efflux pumps but ribosome protections (38) it is logical to assume that it would not be able to provide cross-resistance to CHX. Nevertheless, further experiments are required to confirm our hypotheses.

Our findings constitute an example of the discovery of a new resistance mechanism in *Enterobacteriaceae* through GWAS. Here such analysis using phenotypic and genomic data allowed to identify TetA of class B as a determinant of CHX resistance. Formerly, such approach had been successfully used to decipher antibiotic resistance in various species of clinical interest as for example *Mycobacterium tuberculosis* (39), *Streptococcus pneumoniae* (40), *Neisseria gonorrhoeae* (41) as well as *Staphylococcus aureus* (42). In addition, GWAS allowed the identification of pivotal determinants of virulence in *E. coli* (43, 44).

In conclusion, we showed that natural environments cannot be reduced to the test tube as evidenced by the two different evolutionary paths leading to reduced susceptibility in *E. coli*. On one hand, mutations in the chromosomal gene *mlaA*, with corresponding protein being involved in retrograde transport of phospholipids, was only observed among *in vitro* mutants and likely associated with a high fitness cost in natural environments (36, 45). On the other hand, a TetA-type efflux mechanism mediates decreased susceptibility to CHX, being highly prevalent among *E. coli* isolates. Studies are needed to identify the primary selective pressure leading to the high frequencies of these strains in intensive care unit particularly in ventilator-assisted pneumonia. It is possible that the spread of *tetA* efflux gene is associated to the spread of *Tn*10 transposons. Those transposons are widespread and easily exchanged among Gram-negative bacteria and their spread is enhanced by tetracycline. It is therefore tempting to speculate that the spread of those *Tn*10 *tetA* associated genes are related to the use of tetracycline in human and veterinary medicines, and the subsequent presence of tetracycline residues in large amount in the *E. coli* secondary habitat, water and sediments (46).

## Acknowledgements

We thank Marie Petitjean for kindly providing us data on antibiotic resistance in EnteroBase.

