## Supplementary figures and images for "Chlorhexidine reduced susceptibility associated to tetracycline resistance in clinical isolates of *Escherichia coli*"

### Figure S1

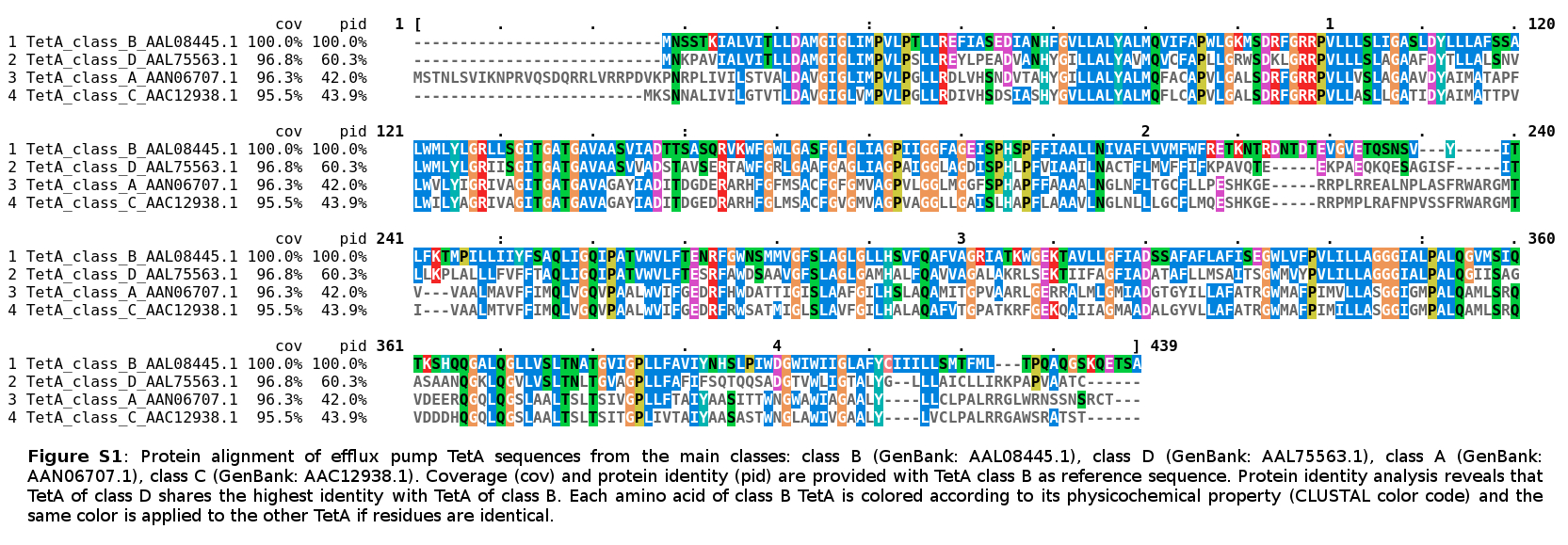
